# Developing a newborn rat model of meningitis without concomitant bacteremia by intraventricular injection of K1 (-) *Escherichia coli*

**DOI:** 10.1101/384750

**Authors:** Yun Sil Chang, So Yoon Ahn, Dong Kyung Sung, Young Eun Kim, Se In Sung, So Yoon Joo, Won Soon Park

## Abstract

Neonatal meningitis caused by *Escherichia coli* results in high mortality and neurological disabilities, and the concomitant systemic bacteremia confounds its mortality and brain injury. This study developed an experimental model of neonatal meningitis without concomitant systemic bacteremia by determining the bacterial inoculum of K1 capsule-negative *E. coli* by intraventricular injection in newborn rats. Meningitis was induced by intraventricular intraventricular injection of 1 × 10^2^ (low dose), 5 × 10^2^ (medium dose), or 1 × 10^3^ (high dose) colony forming units (CFU) of K1 (-) *E. coli* (EC5ME) in Sprague-Dawley rats at postnatal day 11. Ampicillin was started at postnatal day 12. Blood and cerebrospinal fluid (CSF) cultures were performed at 6 h, 1 day, and 6 days after inoculation. Brain magnetic resonance imaging (MRI) was performed at postnatal days 12 and 17. Survival was monitored, and brain tissues were obtained for histological and biochemical analyses at P12 and P17. Survival was inoculum dose-dependent, with lowest survival in high dose group (20&#0x0025;) compared with medium (80%) or low (70%) dose group. CSF bacterial counts in low and medium dose group were significantly lower than that in high dose group at 6 h, but not at 24 h after inoculation. No bacteria were isolated from the blood throughout the experiment, or from the CSF at postnatal day 17. Brain MRI showed an inoculum dose-dependent increase in the extent of ventriculomegaly, cerebral infarct, extent of brain injury, and inflammatory responses. We developed a newborn rat model of bacterial meningitis without concomitant systemic bacteremia by intraventricular injection of K1 (-) *E.coli*.

## Introduction

Despite continuous improvements in antibiotic therapy and intensive care medicine, bacterial meningitis remains a serious disease at any age, and the prognosis is particularly poor in newborn infants, with mortality rates of 20–40% and long-term neurological sequelae, including deafness, blindness, seizures, hydrocephalus, and cognitive impairment in up to 50% of the survivors(1-3). The precise mechanisms by which bacterial infection and the ensuing inflammatory responses in the subarachnoid space during neonatal bacterial meningitis lead to neuronal injury that could result in death or neurological sequelae in survivors are not completely delineated. Therefore, a better understanding of the mechanism of brain damage is necessary to prevent this neuronal injury, and consequently to reduce the mortality and morbidities associated with neonatal bacterial meningitis.

Developing an appropriate animal model that could simulate clinical bacterial meningitis in newborn infants would be essential to determine its pathogenesis, and also to test the efficacy of newly developed adjuvant treatments in addition to the use of antibiotics. Currently, several animal models of neonatal bacterial meningitis, including newborn piglets (4), mice (5-7), rats (8, 9), or rabbits(10) are available, and meningitis was induced by various routes including intraperitoneal (5, 11), intranasal (6), intravenous (5, 10, 12), or intracisternal (7-10, 12) inoculation of bacteria. However, these animal models have certain drawbacks, including small sample size, low infectivity, high mortality, and/or variable extent of brain injury (11). Furthermore, concomitant bacteremia might aggravate the meningitis-induced brain injury (9, 13, 14), thus increasing mortality (8, 9, 15). Therefore, in the present study, we developed a newborn rat model of neonatal bacterial meningitis to mimic the human clinical and neuropathological abnormalities, using 11-day-old newborn Sprague–Dawley rats with titrated intraventricular inoculation of *Escherichia coli*, the most common gram-negative pathogen of neonatal bacterial meningitis (3). We attempted to determine the bacterial inoculum dose with maximal brain injury and minimal mortality by using K1 capsule-negative *E. coli* to confine the infection to the central nervous system, without concomitant systemic bacteremia (12, 16). We inoculated the bacteria intraventricularly using a stereotaxic frame to simulate the neuropathological progression of clinical neonatal bacterial meningitis, which begins with ventriculitis (17, 18). Brain injury was monitored *in vivo* by brain magnetic resonance imaging (MRI) (19-22).

## Materials and Methods

### Infecting organism

We used EC5ME, an un-encapsulated mutant of *E. coli* strain possessing the K1 capsular polysaccharide C5 (serotype 018:K1:H7) (a kind gift from Professor Kwang Sik Kim, Johns Hopkins University, MD, USA)(12, 16) to induce only bacterial meningitis, but not secondary bacteremia, in this study. Bacteria were cultured overnight in brain heart infusion broth, diluted in fresh medium, and grown for another 6 h to mid-logarithmic phase. The culture was centrifuged at 5,000 ×g for 10 min, re-suspended in sterile normal saline to the desired concentration, and used for intraventricular injection. The accuracy of the inoculum size was confirmed by serial dilution, overnight culture on blood agar plates, and then count of colony forming units (CFU).

### Animal model of meningitis

The experimental protocols described herein including anticipated mortality was reviewed and approved by the Animal Care and Use Committee of Samsung Biomedical Research Institute which provides special training in animal care or handling for research staff. All animal procedures were performed in an AAALAC-accredited specific pathogen-free facility and done in accordance with Institutional and National Institutes of Health Guidelines for Laboratory Animal Care. Fig 1 shows details of the experimental schedule. The experiment began at P11, and continued through to P17. We assessed and monitored the condition of rat pups on a daily basis regularly. To induce meningitis, newborn Sprague–Dawley rats (Orient Co, Seoul, Korea) were anesthetized using 2% isoflurane in oxygen enriched air, and a total of 10 μl EC5ME inoculum in saline was slowly infused into the left ventricle under stereotactic guidance (Digital Stereotaxic Instrument with Fine Drive, MyNeurolab, St. Louis, MO, USA; coordinates: x = ± 0.5, y = ± 1.0, z = ± 2.5 mm relative to the bregma) at P11. To determine the optimal inoculum dose with minimal mortality and maximal brain injury, we tested three different inoculum doses of *E. coli:* A low inoculum dose of 1 × 10^2^ CFU EC5ME (LE), a medium inoculum dose of 5 × 10^2^ CFU EC5ME (ME), and a high inoculum dose of 1 × 10^3^ CFU EC5ME (HE). For normal control group (NC), equal volume of normal saline was given intraventricularly. After the procedure, the rat pups were allowed to recover and returned to their dams, and there was no mortality associated with the procedure. First, 10 rat pups for each group were allocated to assess the acute pathophysiological changes, and the survivors were sacrificed at 24 h (P12) after bacterial inoculation for histopathological assessment (n = 6, 5, 4 and 3 for the NC, LE, ME and HE groups, respectively) and biochemical analyses (n = 4, 4, 4 and 3 for the NC, LE, ME and HE groups, respectively). We also conducted the time course experiment in 10 animals for each group to determine the survival rate until sacrifice of the survivors at P17 for histopathological assessment (n = 5, 4, 4 and 2 for the NC, LE, ME and HE groups, respectively) and biochemical analyses (n = 5, 3, 4 and 0 for the NC, LE, ME and HE groups, respectively). Intraperitoneal injection of ampicillin (200 mg/kg/day) was started 6 h after bacterial inoculation, and continued for 3 days until P13. CSF was obtained to determine the bacterial titer at 6 h, 24 h, and 6 days (P17) after bacterial inoculation. Brain MRI was performed at P12 and P17. All experimental procedures generating pain were performed under the isoflurane inhaled anesthesia to reduce pain. All animals were daily monitored and we assessed mortality. Every cause of death was not associated with experimental procedures and related to disease condition. At P12 and P17, survived animals were euthanized by isoflurane and sacrificed by cervical vertebra dislocation and whole brain tissue and CSF samples were obtained.

**Fig 1.**
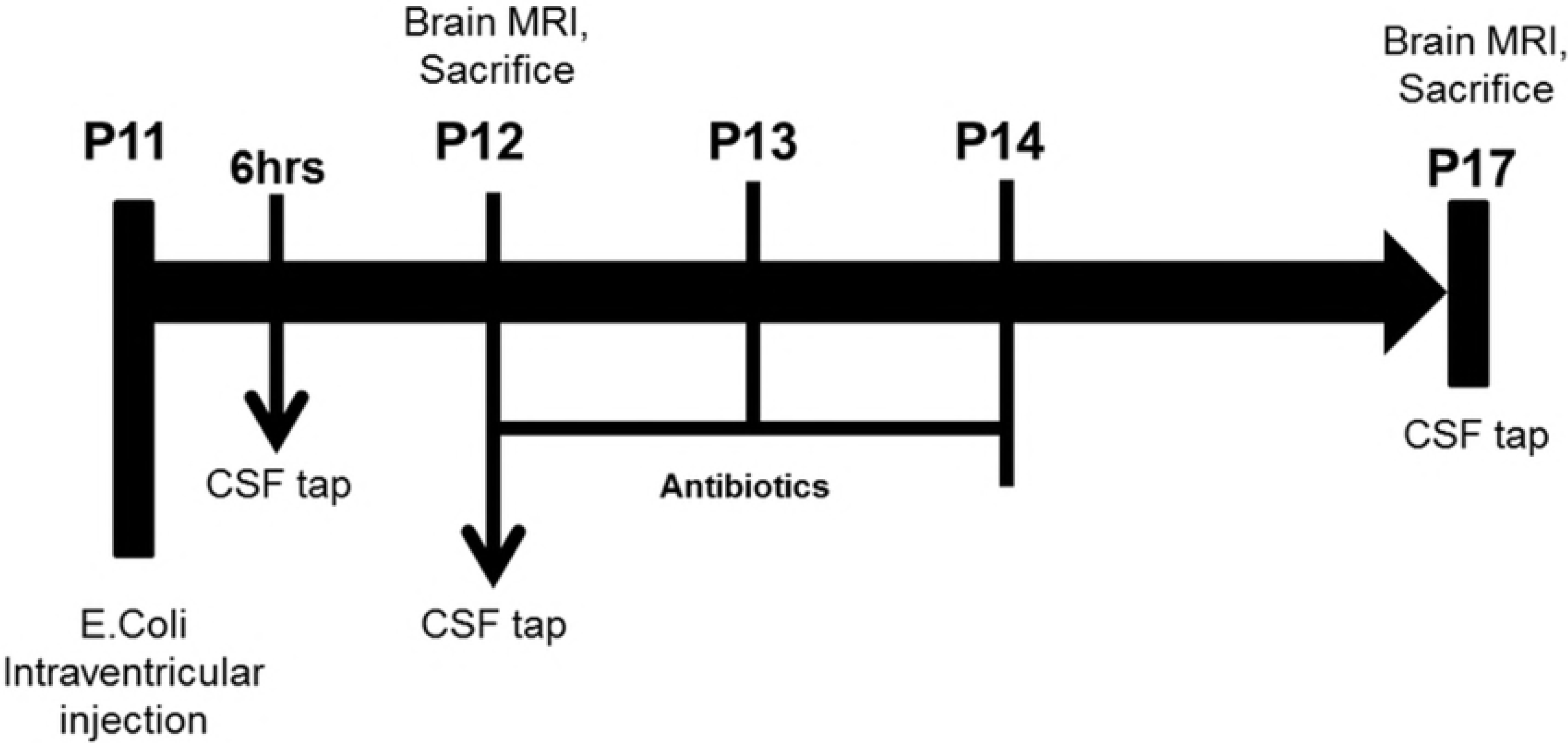
Experimental protocol. *E. coli* was injected intracerebroventriculary on P11 at different doses for each group; low dose of 1 × 10^2^ CFU, a medium dose of 5 × 10^2^ CFU, and a high dose of 1 × 10^3^ CFU. Brain MRI was performed before the rats were sacrificed.

### Bacterial quantification

Bacterial concentrations from each study group were measured in the CSF and blood at 6 h, 24 h, and 6 days after bacterial inoculation for induction of meningitis. Bacteria CFU levels in the CSF and blood were measured at dilutions of 10^−4^–10^−8^ plated on brain heart infusion agar after overnight incubation at 37°C.

### In vivo brain MRI assessment

The brain MRI was performed while the rats were kept in an anesthetized state by the administration of 1.5–2% isoflurane in oxygen-enriched air using a facemask. All MRI examinations were performed using a 7.0-tesla MRI System (Bruker-Biospin, Fällanden, Switzerland) prepared with a 20-cm gradient set capable of providing a rising time of 400 mTm-1. The MR images were acquired with 1.0-mm slice thickness, and a total of 12 slices were acquired. Brain MRI was performed at P12 (n = 10, 9, 8 and 6 in the NC, LE, ME and HE groups, respectively) and at P17 (n = 11, 7, 8 and 2 in the NC, LE, ME and HE groups, respectively). After the MRI exams, the rat pups were allowed to recover and were returned to their dams.

### Measurement of the extent of brain injury by MRI

All MR images were analyzed using Image J software (National Institutes of Health). The lesion was well identified by the hyperintense areas in DWI at P12 and by the hyperintense areas in T2-weighted imaging at P17. The ratio of the infarcted region in the cortex to the whole brain volume was calculated as a parameter of brain injury. The ventriculomegaly volume ratio was also calculated for each pup.

### Tissue preparation

Brain tissue preparation procedures were performed in the surviving animals until P12 (n = 9, 8 and 6 in the NC, LE, ME and HE groups, respectively) and P17 (n = 10, 7, 8 and 2 in the NC, LE, ME and HE groups, respectively). The animals were anesthetized with sodium pentobarbital (100 mg/kg), and their brains were isolated after thoracotomy and transcardiac perfusion with ice-cold 4% paraformaldehyde in 0.1 mol/l phosphate-buffered saline (PBS). The brains were carefully removed from the animals and fixed overnight with 4% formaldehyde solution at room temperature. The brains were embedded in paraffin, and coronal serial sections (4-μm thick) were taken from the paraffin blocks for morphometric analyses at the level of the medial septum area (+0.95 mm to −0.11/bregma) and the hippocampal area (−2.85 to −3.70 mm). The sections were stained with hematoxylin and eosin to assess the extent of neuronal damage.

### TUNEL Assay

Cell death in the hippocampal region was assessed using the immunofluorescent terminal deoxynycleotidyltransferase-mediated deoxyuridine triphosphate nick-end labeling (TUNEL) technique (kit G3250, Promega, Madison, USA). The slides were mounted with Vectashield mounting solution with 4′, 6′-diamidino-2-phenylindole dihydrochloride hydrate (DAPI; H-1200; Vector) and visualized by 20× (dentate gyrus) and 5× tiles can confocal microscopy (Leica, Wetzlar, Germany). A blinded evaluator counted the density of TUNEL-positive nuclei in whole brain on coronal brain sections. Six coronal sections (+0.95 mm to −0.11 mm/bregma) were counted from each brain.

### Immunohistochemistry

Immunohistochemistry of gliosis (neuronal specific glial fibrillary acidic protein [GFAP]) and reactive microglia (ED-1) was performed on deparaffinized 4-μm thick brain sections. The slices were incubated with the primary anti-GFAP antibodies (rabbit polyclonal; Dako, Glostrup, Denmark, overnight, 4 °C, 1:1,000 in PBS with 1% bovine serum albumin) and the anti-ED-1 antibodies (mouse polyclonal; Millipore, CA, USA, overnight, 4 °C, 1:500 in PBS with 1% bovine serum albumin). After three rinses (same buffer), the sections were incubated with Alexa Fluor 568 (red) conjugated anti-rabbit immunoglobulin (90 min, diluted 1:500; Molecular Probes, Eugene, OR, USA) and Alexa Fluor 568 (red) conjugated anti-mouse immunoglobulin (90 min, diluted 1:500; Molecular Probes, Eugene Oregon) each. After three rinses, the sections were mounted with Vectashield mounting solution containing 4′, 6′-diamidino-2-phenylindole dihydrochloride hydrate and visualized by 20× (dentate gyrus) and 5× tilescan confocal microscopy (Leica). The density of GFAP-positive cells and the number of ED-1-positive cells were determined by a blinded observer in whole tilescan fields of each animal’s brain using ImageJ software.

### Enzyme-linked immunosorbent assay (ELISA)

IL-1α, IL-1β, IL-6, and TNF-α concentrations in tissue homogenates were measured at P12 and P17 using the Milliplex MAP ELISA Kit according to the manufacturer’s protocol (Millipore, Billerica, MA, USA).

### Statistical analyses

Statistical analyses were performed using SPSS version 18.0 (IBM, Chicago, IL, USA). Data are expressed as the mean ± standard error of the mean. For continuous variables, statistical comparison between groups was performed using one-way analysis of variance (ANOVA) and Tukey’s post hoc analysis. P < 0.05 was considered statistically significant.

## Results

### Survival rates and body weight

Fig 1 shows the details of the experimental schedule. The experiment began at P11 and continued through to P17. To induce meningitis, at P11, three different doses of E. coli were injected into the cerebroventricles of newborn rats; low inoculum dose of 1 × 102 CFU (colony forming unit) EC5ME (LE), a medium inoculum dose of 5 × 102 CFU EC5ME (ME), and a high inoculum dose of 1 × 103 CFU EC5ME (HE). The survival rate after induction of bacterial meningitis was bacterial inoculum dose-dependent, showing the lowest survival rate up to postnatal day (P)17 of 20% for the high inoculum dose (HE), and 70% and 80% for LE and ME doses, respectively (Fig 2A). While survival rate up to P17 in the HE group was significantly lower compared to that in the no inoculum control (NC), the survival rate of the LE and ME groups was not significantly reduced compared with the NC group.

**Fig 2.**
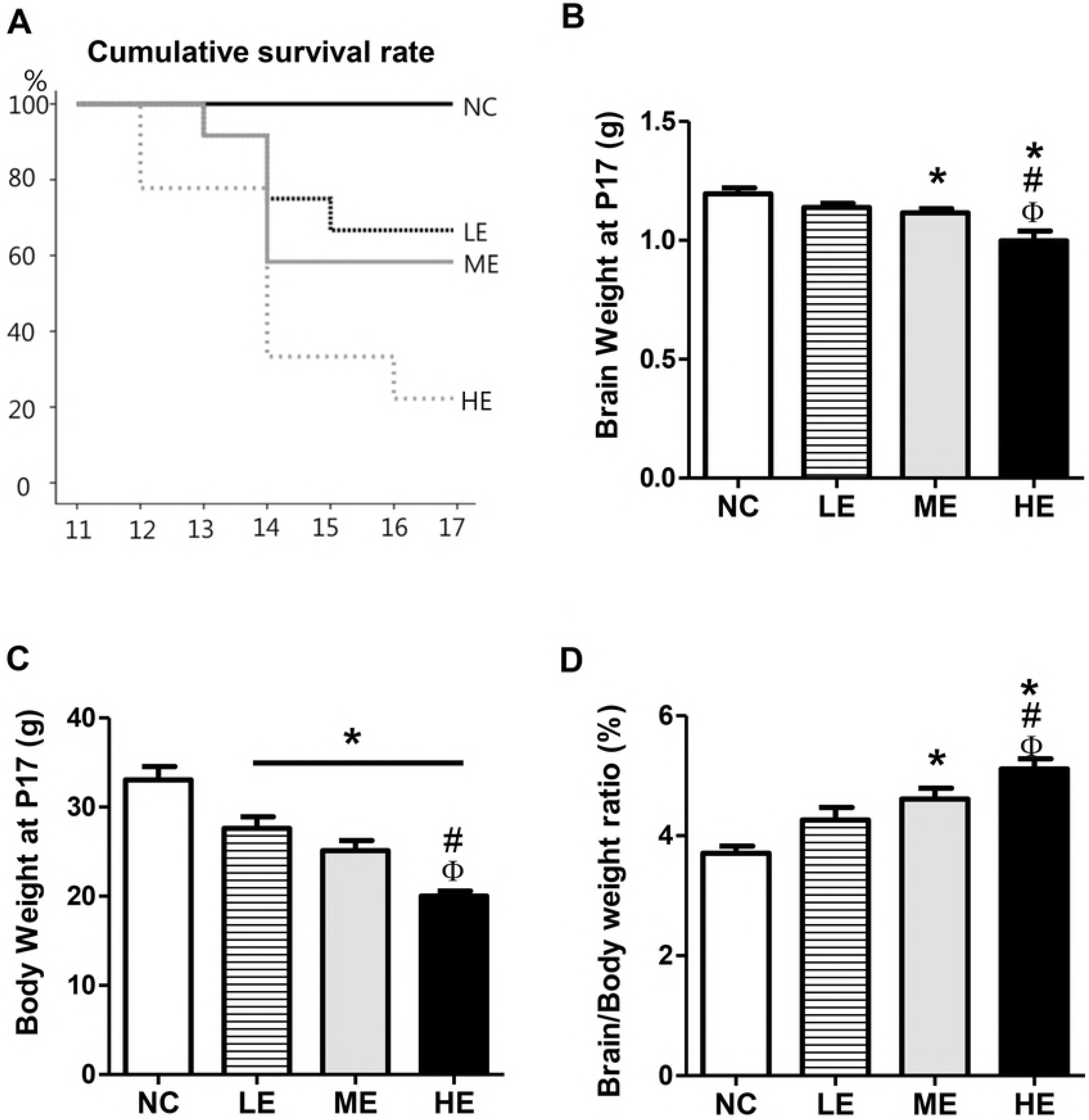
Survival rates. (A) Survival rates in each group were determined using Kaplan–Meier analysis followed by a log-rank test. LE, low dose E. coli group; ME, medium dose E. coli group; HE, high dose E. coli group. (B) Brain weight and (C) body weight were measured at P17 in each group (n=11, 7, 9 and 2 in NC, LE, ME, and HE, respectively). Both weights decreased significantly depending on the E. coli dose. (D) The ratio of brain weight: body weight significantly increased in the HE group compared with the other groups others. Data are presented as the mean ± standard error of the mean (SEM). * P < 0.05 compared with the NC group, # P < 0.05 compared with the LE group, $ P < 0.05 compared with the ME group.

While birth body and brain weight in each study group was not significantly different between the study groups; the body weight gain at P17 in the LE, ME, and HE groups was significantly lower, the brain weight gain in the ME and HE groups was significantly lower, and the brain/body weight ratio in the ME and HE groups was significantly higher compared with the those in the NC group. The least body and brain weight gain, and the highest brain/body ratio, were observed in the HE group compared with those in the LE and ME groups (Fig 2B-D).

### Bacterial counts

To evaluate the bacterial burdens, the CFU were counted in the cerebrospinal fluid (CSF) and blood from each study groups at 6 h (P11), 24 h (P12), and 6 days (P17) after induction of meningitis. While no bacterial growth in the blood was detected in all study groups throughout the experiment, the bacterial counts in the CSF at 6 h after the induction of meningitis in both the LE and ME were significantly lower compare with that in the HE. Thereafter, the bacterial counts in the CSF of all study groups increased significantly compared with that at 6 h, and there were no significant inter-group differences at 24 h after the induction of meningitis (Fig 3). No bacterial growth in the CSF was detected all study groups at 6 days after the induction of meningitis.

**Fig 3.**
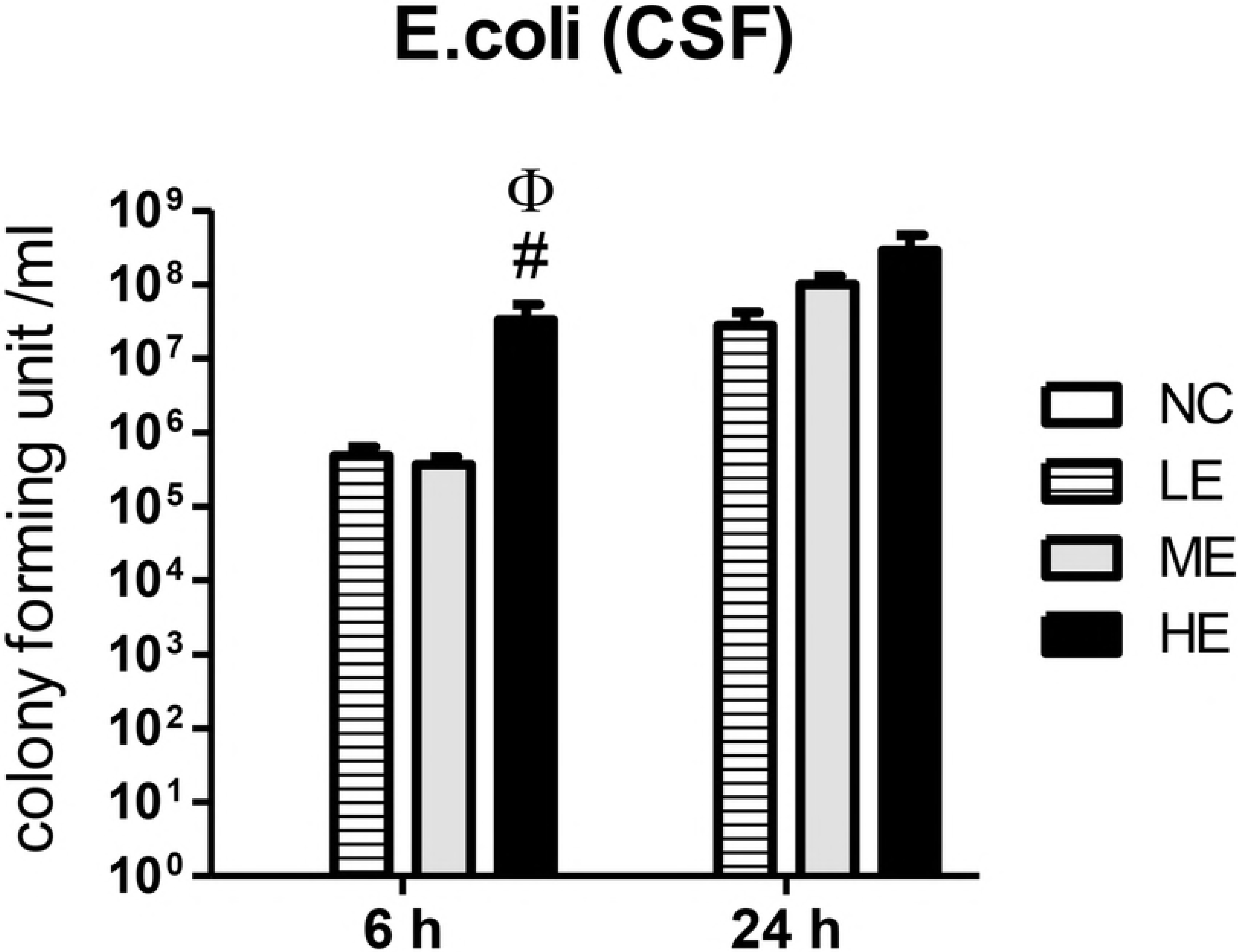
Bacterial counts in the CSF. Bacterial counts in the CSF obtained at 6 and 24 h after bacterial inoculation and before initiation of antibiotic treatment. LE, low dose E. coli group; ME, medium dose E. coli group; HE, high dose E. coli group. Data are presented as the mean ± SEM. * P < 0.05 compared with the NC group, # P < 0.05 compared to LE, $ P < 0.05 compared to ME.

### Brain MRI

To assess the extent of meningitis-induced brain infarction and hydrocephalus, *in vivo* brain MRI scans were taken. The degree of the brain infarct in the ipsilateral cortex and the dilatation of the ventricle to whole brain as evidenced by the hyperintense areas in the diffusion-weighted MRI performed at P12 and by T2-weighted MRI performed at P17 were measured.

The brain infarct volume ratios at P12 and P17 were bacterial inoculum dose-dependently increased, showing the highest ratio in the HE group, and a seemingly increased ratio in the ME group compared with that in the LE group that did not reach statistical significance (Fig 4). The ventriculomegaly volume ratios at P12 were bacterial inoculum dose-dependently increased, showing the highest increase in the HE group compared with that in the LE and ME groups. In addition, although the absolute extent of ventriculomegaly was significantly reduced compared with P12, the ventriculomegaly volume ratios at P17 were also bacterial inoculum dose-dependently increased, showing the highest increase in the HE group compared with that in the LE and ME groups (Fig 4).

**Fig 4.**
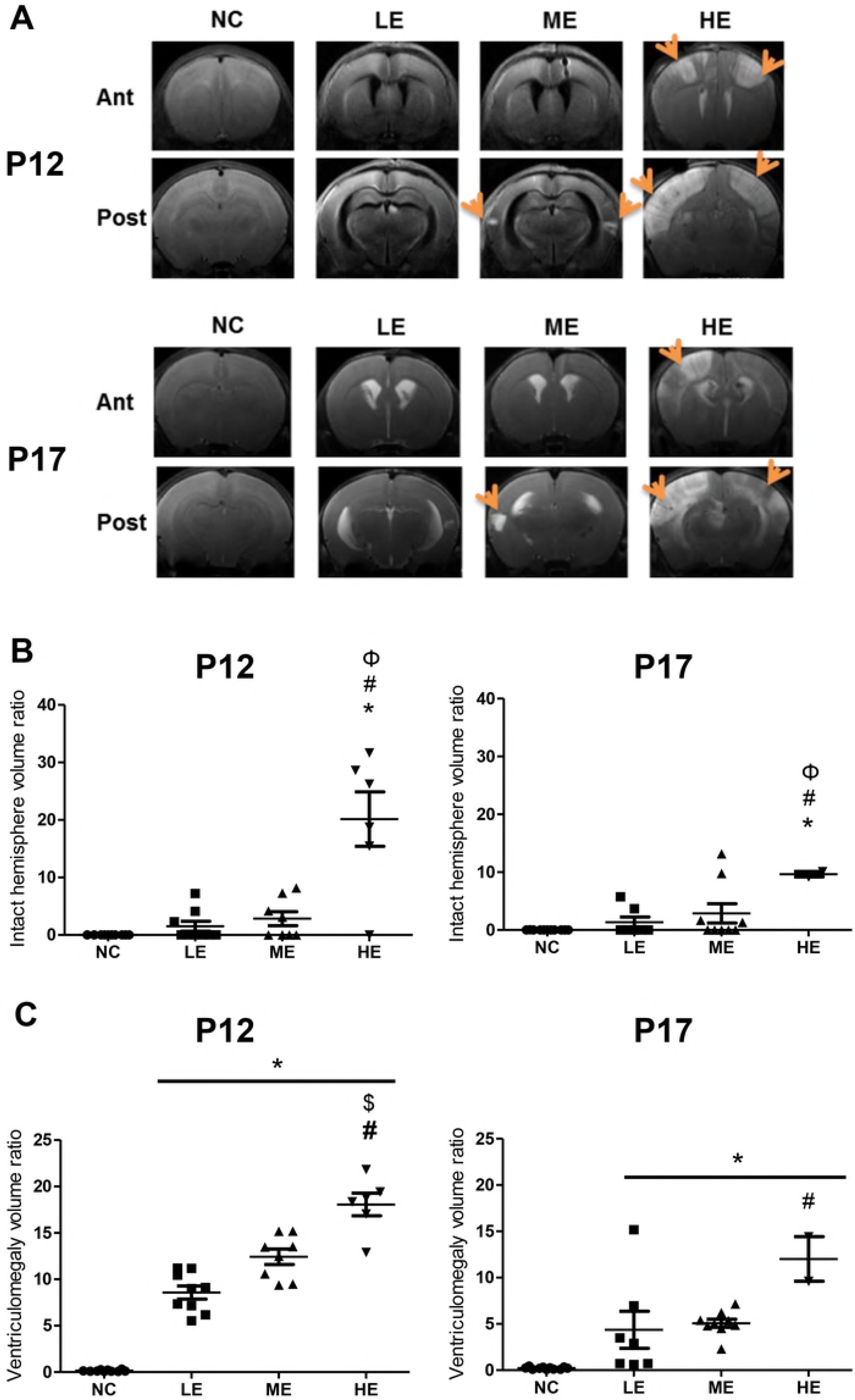
Evolution of brain injury at P12 and P17. (A) Representative brain MRIs of the NE (no *E. coli* control) (left column), LE (middle left column), ME (middle right column), and HE (right column) groups from the medial septal area on day 1 and day 6 after meningitis (P12 and P17). (B) The intact volume of the cortex area to whole brain ratio and (C) the ventriculomegaly volume ratio were measured by MRI at P12 and P17. LE, low dose *E. coli* group; ME, medium dose *E. coli* group; HE, high dose *E. coli* group. Data are presented as the mean ± SEM. * P < 0.05 compared with the NC group, # P < 0.05 compared with the LE group, $ P < 0.05 compared with the ME group.

### TUNEL staining and immunohistochemistry

To assess the extent of bacterial meningitis-induced cell death, and reactivate gliosis and microglia in the brain, the number of terminal deoxynycleotidyltransferase-mediated deoxyuridine triphosphate nick-end labeling (TUNEL)- and ED-1 (Ectodysplasin A) positive cells, and the density of glial fibrillary acidic protein (GFAP)-positive cells in the hippocampus were estimated at 24 h after induction of meningitis (P12). The number of TUNEL- and ED-1 positive cells, and the intensity of GFAP-positive cells in the hippocampus at P12 were bacterial inoculum dose-dependently increased compared with the NC group, showing the highest increase in the HE group. The increased number of TUNEL positive cells and the intensity of GFAP positive cells in the ME group were significantly higher compared with those in the LE group (Fig 5).

**Fig 5.**
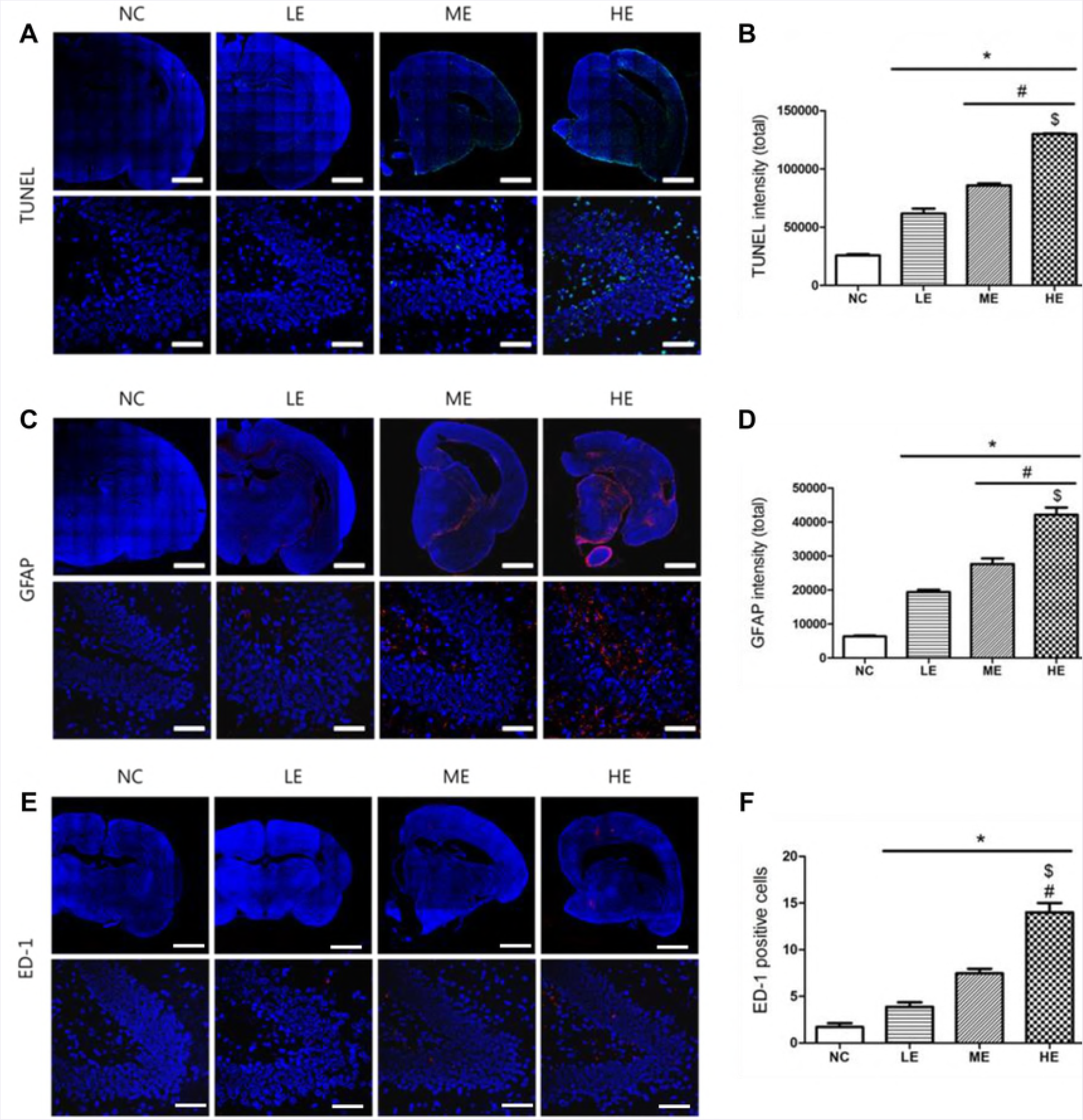
Immunostaining in the hippocampus region. Representative photomicrographs of (A) TUNEL, (C) GFAP intensity, and (E) ED-1 positive cells in the brain of P12 rats in each group. TUNEL intensity was labeled with FITC (green); GFAP and ED-1 positive cells were labeled with TRITC (red). The cell nuclei were labeled with DAPI (blue) (Scale bar = 25 p,m). The average intensity of observed (B) TUNEL and (D) GFAP, and the average number of (F) ED-1 positive cells per high-power field (HPF) in each group are also represented. LE, low dose *E. coli* group; ME, medium dose *E. coli* group; HE, high dose *E. coli* group. Data are presented as the mean ± SEM. * P < 0.05 compared with the NC group, # P < 0.05 compared with the LE group, $ P < 0.05 compared with the ME group.

### Inflammatory Cytokines in Brain

Levels of inflammatory cytokines, such as interleukin (IL)-1α, IL-1β, IL-6, and tumor necrosis factor alpha (TNF-α) measured in the periventricular brain tissue homogenates at P12 revealed bacterial inoculum dose-dependent increase, showing the highest increase in the HE group. The inflammatory cytokine levels in the ME group were significantly higher compared with those in the LE group (Fig 6). Although the brain homogenates of the HE group were not available for measurements because of their high mortality at P17, and the absolute levels of the inflammatory cytokines were significantly reduced compared with P12, the inflammatory cytokines were bacterial inoculum dose-dependently increased, showing significantly higher levels in the ME group compared with those in the LE group.

**Fig 6.**
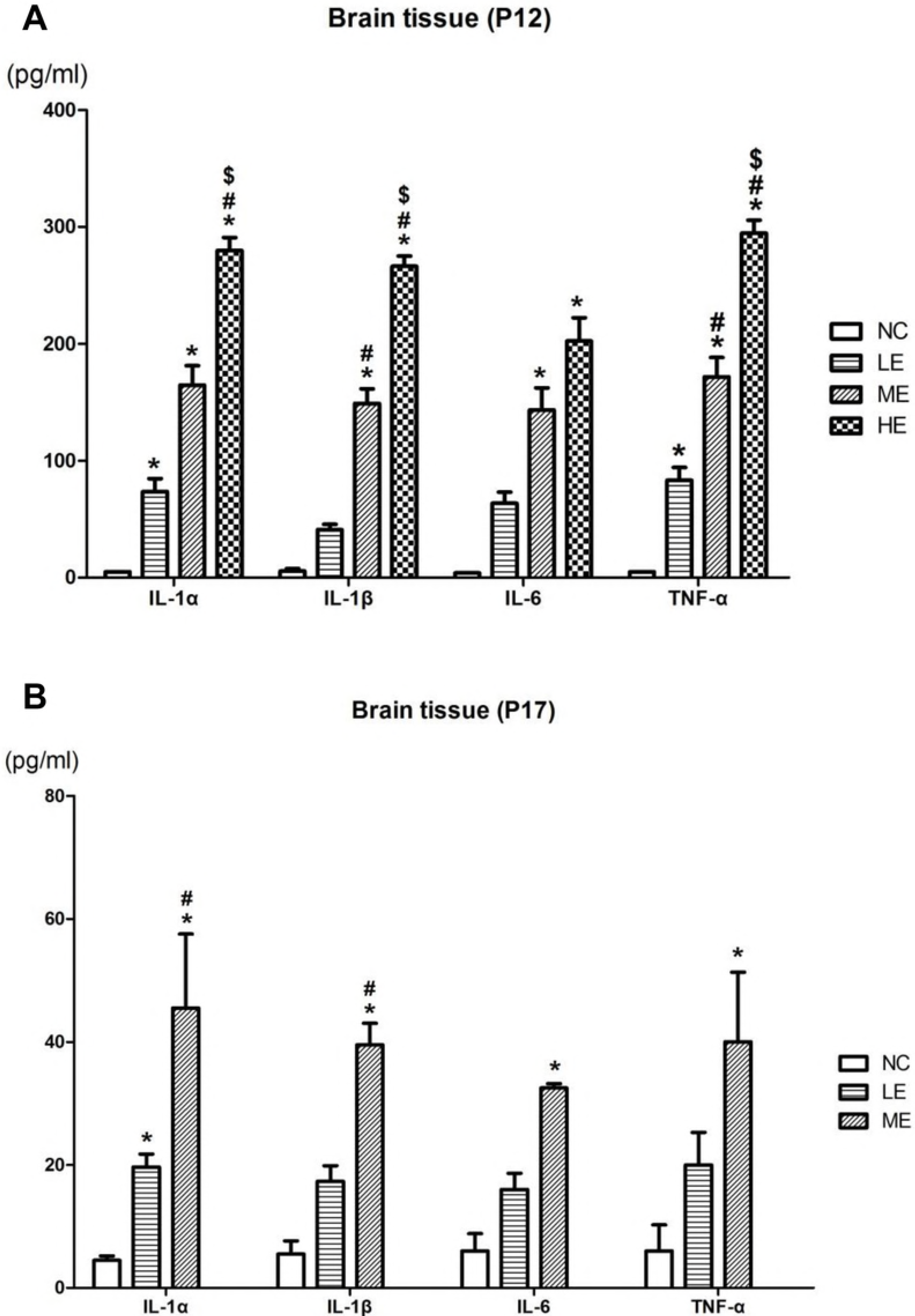
Inflammatory cytokines of brain. > Interleukin [IL]-1α, IL-1β, IL-6, and tumor necrosis factor [TNF]-α concentrations in brain tissue homogenates at (A) P12 and (B) P17, were measured using ELISA in each group. LE, low dose *E. coli* group; ME, medium dose *E. coli* group; HE, high dose *E. coli* group. Data are presented as the mean ± SEM. * P < 0.05 compared with the NC group, # P < 0.05 compared with the LE group, $ P < 0.05 compared with the ME group.

## Discussion

Despite recent improvements in neonatal intensive care medicine and development of highly active new antibiotics, neonatal bacterial meningitis remains a serious disease with high mortality and neurological morbidities in survivors(1, 3). Currently, few effective adjuvant therapies are available to improve the prognosis of this intractable and devastating neonatal disorder. Therefore, developing an appropriate animal model to simulate clinical bacterial meningitis in newborn infants is an essential first step to determine its pathophysiological mechanisms, and to test the therapeutic efficacy of any potential new treatments. However, the limitations of currently available experimental models of meningitis lie in the great variability between the species, the inoculation methods, and the age of the animal models (11). In this study, we used P11 rats as an animal model of neonatal meningitis because the rat brain at P11 is comparable in terms of maturation to the human brain at birth (23). The larger size of rat pups compared with mice enables easier surgical manipulation at an earlier age, and a larger amount of brain tissues obtained at harvest. Furthermore, our already established newborn rat model of severe Intraventricular hemorrhage (20-22), middle cerebral arterial occlusion (24), and hypoxic ischemic encephalopathy (25) with *in vivo* brain MRI and histopathological analyses to study the pathophysiological mechanisms and therapeutic efficacy could be easily extrapolated to develop a newborn rat model of meningitis in this study. Overall, the findings of the present study suggested that the newborn rat pup model is suitable and appropriate to research the pathogenesis of neonatal bacterial meningitis and to test the efficacy new treatments.

In this study, *E. coli* was used to induce meningitis, because it is the most frequent gram-negative pathogen of neonatal bacterial meningitis (3). Although brain injury primarily results from local meningeal infection, concomitant systemic bacteremia might aggravate the meningitis-induced disease severity, brain injury, and mortality (9, 13-15). This discordance between disease severity and brain injury means that a poorer outcome does not necessarily lead to increased brain injury (15, 19). In addition, neuroprotection might not be associated with improved clinical status (26). Therefore, developing an animal model of neonatal meningitis that could dissect the role of local meningeal infection and systemic bacteremia is essential to evaluate the pathophysiological mechanism of brain injury and to test the therapeutic efficacy of any new treatment approaches to reduce the meningitis-induced sequelae and to improve outcome and survival. As the K1 capsule is the critical determinant for developing *E. coli* meningitis in rats, we used K1(-) *E. coli* in this study to prevent secondary systemic bacteremia (12, 16). Although we observed secondary bacterial invasion from the central nervous system (CNS) into the blood stream once the bacterial concentration in the CSF reached above 10^5^ CFU in our previous experimental study of *E. coli* meningitis in newborn piglets (27), in the present study, we observed no concomitant secondary bacteremia, despite high bacterial concentrations in the CSF well above 10^5^ CFU. Overall, the use of K1 (-) *E. coli* is suitable to study the pathophysiological consequences of meningitis only and the effects of various therapeutic interventions, without the confounding effects of simultaneous systemic bacteremia.

The neuropathology of neonatal bacterial meningitis begins with choroid plexitis and ventriculitis (18, 28, 29), and progresses to arachnoditis and vasculitis, leading to brain edema, hydrocephalus, infarction, and periventricular leukomalacia (30). In the present study, *K1 (-) E. coli* was injected intraventricularly to induce meningitis because although it bypasses the natural hematogenous bacterial invasion across the blood brain barrier into the CNS(12, 16), this experimental model is more clinically relevant by simulating the clinical neuropathological progression of neonatal bacterial meningitis beginning with ventriculitis (18, 28, 29).

In the present study, we tested three different doses of K1 (-) *E. coli* (EC5ME) for the induction of meningitis to determine the optimal inoculum dose with minimal mortality and maximal brain injury; 1 × 10^2^ CFU for the LE group, 5 × 10^2^ CFU for the ME group, and 1 × 10^3^ CFUs for the HE group. Survival rates, body and brain weight gain, the extent of inflammatory responses and brain injury correlated significantly with the inoculum dose used to induce meningitis, showing highest mortality, extent of inflammatory responses, and brain injury, and the least body and brain weight gain. We also observed higher inflammatory responses and the least extent of brain injury in the ME and LE groups, respectively. The mortality rate was positively correlated with the inoculum dose and the extent of inflammatory responses and brain injury. As blood culture was negative throughout the experiment, the inoculum dose-dependent increase in mortality, inflammatory responses, and brain injury solely reflects the virulence of EC5ME meningitis, without the confounding effects of the concomitant systemic bacteremia. Overall, these findings suggest that ME (5 × 10^2^ CFU) of EC5ME might be optimal inoculum dose to induce neonatal meningitis.

Because bacterial meningitis induces high mortality in newborn infants, the design of animal study was also driven to target for severe, end-stage models. From an ethics viewpoint, this use contradicts views that death as an endpoint is unacceptable. However, the use of alternative end point can generate scientific concerns. Because minor improvements in mortality rates are regarded as major advances in treatment, indefinite endpoint may skew data. Thus, for the development of neonatal meningitis model with proper mortality, precise mortality rate without premature euthanasia was required. Unfortunately, in meningitis model, replacement of animal model is extremely difficult because *in vivo* immune response is too complicated to model in *in vitro* system. For the animal welfare, the development of appropriate model which we aimed in the present study would be essential to reduce animal numbers and may be the most valuable refinement for meningitis study.

In infants with bacterial meningitis, brain MRI scans showed abnormalities including cerebral infarct, subdural empyema, cerebritis, and hydrocephalus(19). Increased brain ventriculomegaly in the acute phase of bacterial meningitis in adults was associated with increased mortality (31). In agreement with the clinical findings(19, 31), an acute inoculum dose-dependent increase in ventriculomegaly and cerebral infarct was observed at 1 day after the induction of meningitis. In addition, although a less absolute extent of ventriculomegaly and a higher extent of cerebral infarct were observed compared with post-inoculation day 1, the inoculum dose-dependent abnormalities persisted at 6 days after the induction of meningitis. Taken together, these findings suggested that brain MRI could be an early prognostic indicator that would be useful to identify patients requiring further therapeutic interventions, and to assess the therapeutic efficacy of any new treatments, both in clinical and experimental settings of meningitis (19, 31).

In infants with bacterial meningitis, brain MRI scans showed abnormalities including cerebral infarct, subdural empyema, cerebritis, and hydrocephalus(19). Increased brain ventriculomegaly in the acute phase of bacterial meningitis in adults was associated with increased mortality (31). In agreement with the clinical findings(19, 31), an acute inoculum dose-dependent increase in ventriculomegaly and cerebral infarct was observed at 1 day after the induction of meningitis. In addition, although a less absolute extent of ventriculomegaly and a higher extent of cerebral infarct were observed compared with post-inoculation day 1, the inoculum dose-dependent abnormalities persisted at 6 days after the induction of meningitis. Taken together, these findings suggested that brain MRI could be an early prognostic indicator that would be useful to identify patients requiring further therapeutic interventions, and to assess the therapeutic efficacy of any new treatments, both in clinical and experimental settings of meningitis (19, 31).

Brain injuries observed in experimental models of neonatal meningitis are unique in consistently reproducing both hippocampal damage and cortical necrosis (7-9). Inflammatory responses are primarily responsible for the ensuing brain injury in bacterial meningitis (3, 7, 16). In the present study, the extent of inflammatory responses both at post-inoculation day 1 and 6, and the increased number of TUNEL, GFAP, and ED-1 positive cells in the hippocampus at 1 day after induction of meningitis, were associated with the bacterial inoculum dose. Antibiotic treatment was started 24 h after bacterial inoculation, and continued for 3 days: no bacteria were isolated, even in the CSF, at 5 days after the induction of meningitis. Taken together, these findings suggested that increased inflammatory responses, but not increased bacterial proliferation and dissemination, triggered by a higher bacterial inoculum, are primarily responsible for the ensuing brain injury.

In summary, we successfully developed a newborn rat model of neonatal bacterial meningitis without concomitant systemic bacteremia by intraventricular injection of K1 capsule-negative *E. coli* at P11. We also determined that a bacterial inoculum dose of 5 × 10^2^ CFU of EC5ME had the minimum mortality, and maximal inflammatory responses and ensuing brain injury. This animal model is more clinically relevant because neonatal meningitis begins with ventriculitis (18, 28, 29), and could provide the basis for both pathophysiology and intervention studies for neonatal bacterial meningitis not confounded by simultaneous systemic bacteremia. Hopefully, our newly developed newborn rat model of neonatal meningitis will lead to more detailed knowledge of, and new treatments for, this intractable and devastating disorder.

## Author contributions

Yun Sil Chang and So Yoon Ahn contributed equally as co-first authors in conceptualization of the study design and hypothesis, data collection and analysis, manuscript writing and revision. Won Soon Park contributed the study idea, design, and hypothesis, data collection and analysis, critically reviewed and revised the manuscript, and serves as the corresponding author. So Yoon Joo, Dong Kyung Sung, and Young Eun Kim contributed conceptualization of the study design, biochemical analysis and wrote a portion of the manuscript, and critically reviewed and revised the manuscript. All authors listed above have read and approved the manuscript.

## References

1. Anderson SG, Gilbert GL. Neonatal gram negative meningitis: a 10-year review, with reference to outcome and relapse of infection. Journal of paediatrics and child health. 1990;26(4):212–6.

2. Edwards MS, Rench MA, Haffar AA, Murphy MA, Desmond MM, Baker CJ. Long-term sequelae of group B streptococcal meningitis in infants. The Journal of pediatrics. 1985;106(5):717–22.

3. Polin RA, Harris MC. Neonatal bacterial meningitis. Seminars in neonatology: SN. 2001;6(2):157–72.

4. Park WS, Chang YS, Lee M. Effect of induced hyperglycemia on brain cell membrane function and energy metabolism during the early phase of experimental meningitis in newborn piglets. Brain research. 1998;798(1-2):195–203.

5. Tsao N, Chang WW, Liu CC, Lei HY. Development of hematogenous pneumococcal meningitis in adult mice: the role of TNF-alpha. FEMS immunology and medical microbiology. 2002;32(2):133–40.

6. Zwijnenburg PJ, van der Poll T, Florquin S, van Deventer SJ, Roord JJ, van Furth AM. Experimental pneumococcal meningitis in mice: a model of intranasal infection. The Journal of infectious diseases. 2001;183(7):1143–6.

7. Grandgirard D, Steiner O, Tauber MG, Leib SL. An infant mouse model of brain damage in pneumococcal meningitis. Acta neuropathologica. 2007;114(6):609–17.

8. Gianinazzi C, Grandgirard D, Imboden H, Egger L, Meli DN, Bifrare YD, et al. Caspase-3 mediates hippocampal apoptosis in pneumococcal meningitis. Acta neuropathologica. 2003;105(5):499–507.

9. Brandt CT, Holm D, Liptrot M, Ostergaard C, Lundgren JD, Frimodt-Moller N, et al. Impact of bacteremia on the pathogenesis of experimental pneumococcal meningitis. The Journal of infectious diseases. 2008;197(2):235–44.

10. Tauber MG, Sande MA. Pathogenesis of bacterial meningitis: contributions by experimental models in rabbits. Infection. 1984;12 Suppl 1:S3–10.

11. Chiavolini D, Pozzi G, Ricci S. Animal models of Streptococcus pneumoniae disease. Clinical microbiology reviews. 2008;21(4):666–85.

12. Kim KS, Itabashi H, Gemski P, Sadoff J, Warren RL, Cross AS. The K1 capsule is the critical determinant in the development of Escherichia coli meningitis in the rat. The Journal of clinical investigation. 1992;90(3):897–905.

13. Holler JG, Brandt CT, Leib SL, Rowland IJ, Ostergaard C. Increase in hippocampal water diffusion and volume during experimental pneumococcal meningitis is aggravated by bacteremia. BMC infectious diseases. 2014;14:240.

14. Ostergaard C, Leib SL, Rowland I, Brandt CT. Bacteremia causes hippocampal apoptosis in experimental pneumococcal meningitis. BMC infectious diseases. 2010;10:1.

15. Brandt CT, Lundgren JD, Frimodt-Moller N, Christensen T, Benfield T, Espersen F, et al. Blocking of leukocyte accumulation in the cerebrospinal fluid augments bacteremia and increases lethality in experimental pneumococcal meningitis. Journal of neuroimmunology. 2005;166(1-2):126–31.

16. Kim KS. Pathogenesis of bacterial meningitis: from bacteraemia to neuronal injury. Nature reviews Neuroscience. 2003;4(5):376–85.

17. Chua C. Neonatal meningitis and ventriculitis. Journal of the National Medical Association. 1978;70(11):794–5.

18. Berman PH, Banker BQ. Neonatal meningitis. A clinical and pathological study of 29 cases. Pediatrics. 1966;38(1):6–24.

19. Brandt CT, Simonsen H, Liptrot M, Sogaard LV, Lundgren JD, Ostergaard C, et al. In vivo study of experimental pneumococcal meningitis using magnetic resonance imaging. BMC medical imaging. 2008;8:1.

20. Ahn SY, Chang YS, Sung DK, Sung SI, Yoo HS, Lee JH, et al. Mesenchymal stem cells prevent hydrocephalus after severe intraventricular hemorrhage. Stroke; a journal of cerebral circulation. 2013;44(2):497–504.

21. Ahn SY, Chang YS, Sung DK, Sung SI, Ahn JY, Park WS. Pivotal Role of Brain-Derived Neurotrophic Factor Secreted by Mesenchymal Stem Cells in Severe Intraventricular Hemorrhage in Newborn Rats. Cell Transplant. 2017;26(1):145–56.

22. Ahn SY, Chang YS, Sung DK, Sung SI, Yoo HS, Im GH, et al. Optimal Route for Mesenchymal Stem Cells Transplantation after Severe Intraventricular Hemorrhage in Newborn Rats. PloS one. 2015;10(7):e0132919.

23. Romijn HJ, Hofman MA, Gramsbergen A. At what age is the developing cerebral cortex of the rat comparable to that of the full-term newborn human baby? Early human development. 1991;26(1):61–7.

24. Kim ES, Ahn SY, Im GH, Sung DK, Park YR, Choi SH, et al. Human umbilical cord blood-derived mesenchymal stem cell transplantation attenuates severe brain injury by permanent middle cerebral artery occlusion in newborn rats. Pediatric research. 2012;72(3):277–84.

25. Park WS, Sung SI, Ahn SY, Yoo HS, Sung DK, Im GH, et al. Hypothermia augments neuroprotective activity of mesenchymal stem cells for neonatal hypoxic-ischemic encephalopathy. PloS one. 2015;10(3):e0120893.

26. Leib SL, Kim YS, Ferriero DM, Tauber MG. Neuroprotective effect of excitatory amino acid antagonist kynurenic acid in experimental bacterial meningitis. The Journal of infectious diseases. 1996;173(1):166–71.

27. Park WS, Chang YS, Ko SY, Kang MJ, Han JM, Lee M. Effects of microbial invasion on cerebral hemodynamics and oxygenation monitored by near infrared spectroscopy in experimental Escherichia coli meningitis in the newborn piglet. Neurological research. 1999;21(4):391–8.

28. Daum RS, Scheifele DW, Syriopoulou VP, Averill D, Smith AL. Ventricular involvement in experimental Hemophilus influenzae meningitis. The Journal of pediatrics. 1978;93(6):927–30.

29. Gilles FH, Jammes JL, Berenberg W. Neonatal meningitis. The ventricle as a bacterial reservoir. Archives of neurology. 1977;34(9):560–2.

30. Friede RL. Cerebral infarcts complicating neonatal leptomeningitis. Acute and residual lesions. Acta neuropathologica. 1973;23(3):245–53.

31. Sporrborn JL, Knudsen GB, Solling M, Seieroe K, Farre A, Lindhardt BO, et al. Brain ventricular dimensions and relationship to outcome in adult patients with bacterial meningitis. BMC infectious diseases. 2015;15:367.

